# FLASH irradiation enhances the therapeutic index of abdominal radiotherapy for the treatment of ovarian cancer

**DOI:** 10.1101/2019.12.12.873414

**Authors:** Karen Levy, Suchitra Natarajan, Jinghui Wang, Stephanie Chow, Joshua T. Eggold, Phoebe Loo, Rakesh Manjappa, Frederick M. Lartey, Emil Schüler, Lawrie Skinner, Marjan Rafat, Ryan Ko, Anna Kim, Duaa Al Rawi, Rie von Eyben, Oliver Dorigo, Kerriann M. Casey, Edward E. Graves, Karl Bush, Amy S. Yu, Albert C. Koong, Peter G. Maxim, Billy W. Loo, Erinn B. Rankin

## Abstract

Peritoneal metastases are the leading cause of morbidity and mortality in ovarian cancer. Despite current surgery, chemotherapy and targeted therapies, the majority of patients diagnosed with advanced epithelial ovarian cancer develop recurrent disease and overall survival rates remain poor. It is known that ovarian cancer is a radiosensitive tumor. Historically, total abdominal irradiation (TAI) was used as an effective postsurgical adjuvant therapy in the management of chemotherapy sensitive and resistant ovarian cancer. However, TAI fell out of favor due to high toxicity, particularly of the gastrointestinal tract. We have developed a preclinical irradiation platform that allows for total abdominal ultrahigh dose rate FLASH irradiation. We demonstrate that TAI-FLASH reduces radiation-induced intestinal injury in both healthy and tumor-bearing mice compared to conventional dose rate (CONV) irradiation. Single high dose TAI-FLASH reduced mortality from gastrointestinal syndrome, spared gut function and epithelial integrity, and decreased cell death in crypt base columnar cells. Importantly, FLASH and CONV irradiation had similar efficacy in the reduction of ovarian cancer peritoneal metastases. These findings suggest that FLASH irradiation may be an effective strategy to enhance the therapeutic index of radiotherapy for the treatment of metastatic ovarian cancer in women.

## Introduction

Epithelial ovarian cancer (EOC) is the leading cause of gynecological cancer-related deaths among women in developed countries. Mortality rates are particularly high in EOC because the majority of women are diagnosed with advanced stage disease in which the tumor has disseminated beyond the ovaries and pelvic organs to the peritoneum and abdominal organs including the diaphragm, stomach, omentum, liver, and intestines. Despite current surgery, chemotherapy and targeted therapies, 80% of women diagnosed with advanced EOC cancer develop recurrent disease and only 30% of patients survive 5 years following diagnosis ^1^. Currently when compared to conventional cytotoxic chemotherapy alone, there are no approved therapies that significantly improve overall survival rates in women with metastatic ovarian cancer.

Radiation therapy (RT) is the most effective cytotoxic cancer therapy available for the treatment of localized tumors ^2^. Historically, total abdominal irradiation (TAI) was used as an effective postsurgical adjuvant therapy in the management of both chemotherapy sensitive and resistant ovarian cancer ^3-5^. However, given the risk of gastrointestinal and hematopoietic toxicity associated with large field abdominal radiation, chemotherapy has been preferred over abdominal irradiation in the treatment of metastatic ovarian cancer ^6-8^. More recently, there is renewed interest in using radiotherapy to increase the efficacy of chemotherapy, immunotherapy and targeted therapies in the treatment of ovarian cancer ^9^. However, the toxicities associated with abdominal irradiation remain significant concerns.

We and others have recently reported that ultra-high dose rate FLASH irradiation reduces radiation-induced toxicity to multiple normal tissues. In contrast to clinically used conventional dose rates of about 2-20 Gy/minute (CONV), FLASH irradiation at dose rates of >40 Gy/second (more than two orders of magnitude higher) produces less radiation-induced lung fibrosis and radiation-induced neurocognitive impairment after lung and brain irradiation, respectively ^10-12^. FLASH mediated sparing of the skin from radiation-induced necrosis has been demonstrated in both minipigs and cats ^13^. Meanwhile, in preclinical models of lung cancer FLASH achieves similar tumor control as CONV RT ^10^. These findings demonstrated localized FLASH irradiation of the lung, skin or brain could reduce radiation induced toxicities. However, ovarian cancer presents with widespread metastasis throughout the peritoneal cavity and abdomen requiring a wide abdominal irradiation field that includes the highly radiation sensitive intestine. A radiation treatment modality that could reduce gastrointestinal toxicities associated with total abdominal irradiation, while maintaining tumor control, could be transformative in the treatment of ovarian cancer.

Here we demonstrate that total abdominal FLASH RT reduces radiation-induced intestinal injury and preserves intestinal function in both healthy and tumor-bearing mice. Compared to conventional dose rate RT, FLASH RT reduces cell death in intestinal crypt base columnar cells (CBCs) and enhances crypt regeneration. Importantly, FLASH RT provides similar efficacy to CONV in controlling ovarian cancer peritoneal metastases. Thus, our findings identify a potential strategy to enhance the therapeutic index of abdominal irradiation for metastatic ovarian cancer.

## Results

### FLASH irradiation produces less lethality from radiation-induced gastrointestinal syndrome than CONV

To investigate the safety of abdominal FLASH RT, we first examined the effect of abdominal FLASH RT in irradiation-induced lethal injury. For this purpose, we developed a mouse jig with an irradiation field that extends 3 cm in the cranial/caudal direction starting at the 10^th^ rib allowing for abdominal treatment while sparing the pelvic region **(Figure 1A)**. We utilized a clinical linear accelerator modified to generate a 16 MeV electron beam that delivers both uniform treatment across the mouse and a homogenous depth dose through the mouse (1.0 cm) in both FLASH (216 Gy/s) and conventional mode (0.079 Gy/s) **(Supplemental Figure 1A-F)**. We then monitored the survival (time to morbidity) of female C57BL/6 mice treated with 16 Gy abdominal irradiation. All mice treated with conventional irradiation lost more than 25% of their body weight and died or required euthanasia by day 10 post-irradiation **(Figure 1B, Supplementary Figure 1G)**. In contrast, mice treated with 16 Gy FLASH abdominal irradiation had a nadir in body weight at day 6 losing 25% of body weight and 90% of mice recovered and survived more than 90 days post-irradiation **(Figure 1B, Supplementary Figure 1G)**. Complete blood count analysis 96 h post-irradiation demonstrated no significant hematopoietic toxicity after CONV or FLASH **(Supplementary Figure 1H)**, indicating that the observed toxicity was not attributable to bone marrow suppression. Histologic analysis of the jejunum revealed that while all mice exhibited radiation-induced mucosal damage, the FLASH treated mice had a 2-fold increase in the presence of regenerating crypts at 96 h post-irradiation **(Figure 1C-D)**. Notably, the intestinal mucosa of surviving 16 Gy FLASH treated mice was histologically indistinguishable from unirradiated control animals at 12 weeks post irradiation **(Figure 1E)**. These findings demonstrate that FLASH irradiation produces less radiation-induced lethal injury.

**Figure 1:**
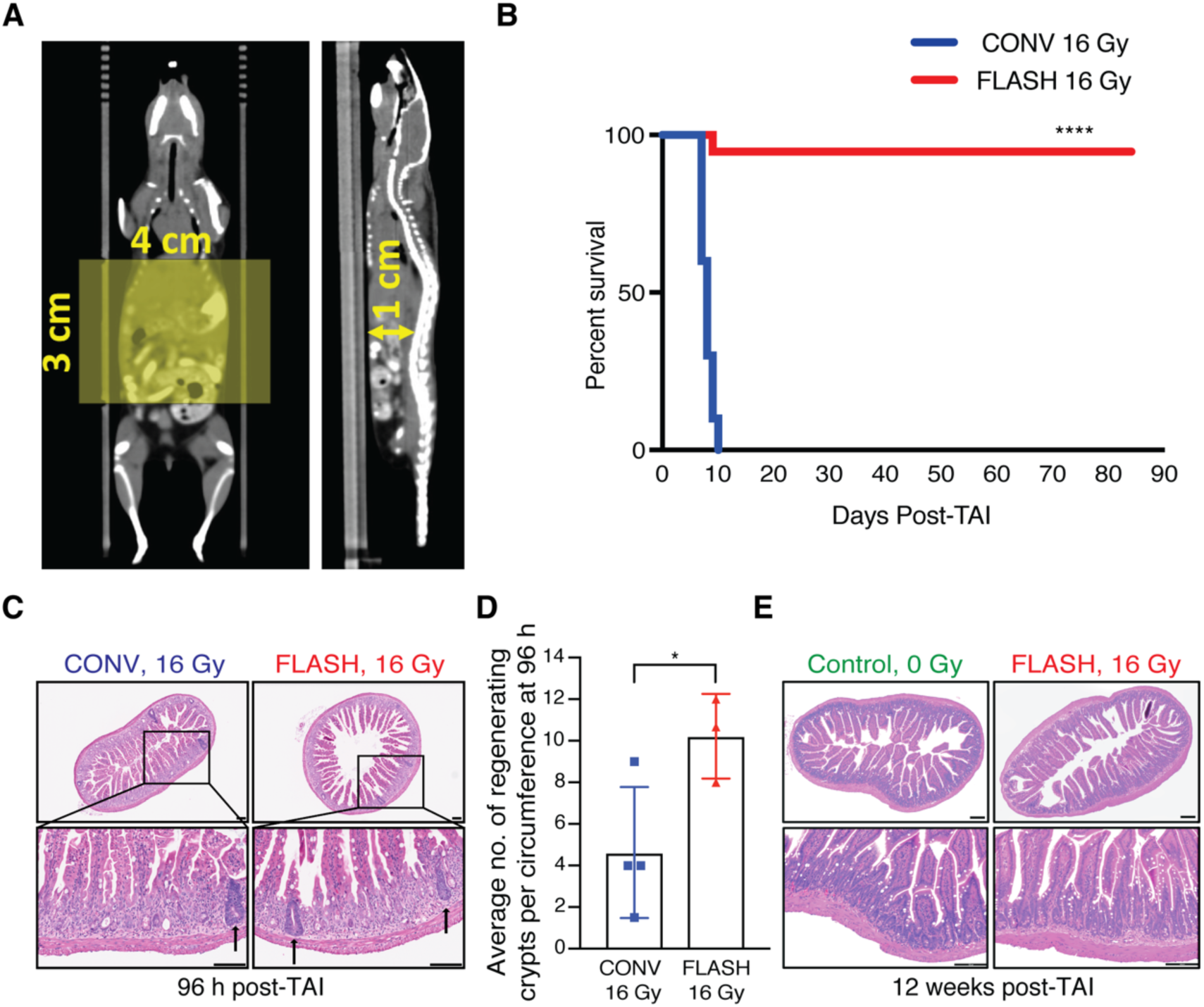
FLASH produces less lethality from radiation-induced gastrointestinal syndrome than CONV irradiation in non-tumor bearing mice. **(A)** CT images (coronal and sagittal slices) showing reproducible mouse positioning within the stereotactic frame. The yellow shaded rectangle (3 cm × 4 cm) indicates the irradiated region in the abdomen; the 3 cm length with the cranial border at the 10^th^ rib encompasses the entire small bowel. The distance from the abdominal wall to the ventral surface of the spine, representing the most posterior extent of the abdominal cavity, is 1 cm (yellow arrow). **(B)** Kaplan-Meier survival curve of animals that received 16 Gy total abdominal irradiation (TAI), demonstrating complete lethality after CONV but near-complete survival after FLASH irradiation (n=10 per group, **** by log-rank (Mantel-Cox) test p<0.0001). **(C)** Histological images of hematoxylin and eosin (H&E) stained jejunal sections from animals 96 h after 16 Gy TAI. Arrows point to regenerating crypts. **(D)** Quantification of the average number of regenerating crypts per circumference 96 h post 16 Gy TAI, demonstrating over double the number of regenerating crypts after FLASH vs. CONV irradiation. Regenerating crypts were counted in 3 mid-jejunal circumferences per mouse (n=4 mice in CONV; n=3 in FLASH, *p<0.05 by unpaired 2-tailed Student’s t-test). **(E)** Histological images of H&E stained jejunum cross-sections from unirradiated mice and surviving mice 12 weeks after 16 Gy FLASH TAI, demonstrating normal histology after recovery from FLASH. Scale bar: 100 μm. Error bars represent standard deviation of the mean.

### FLASH irradiation spares intestinal function and epithelial integrity compared to CONV

We next evaluated a sublethal dose of irradiation (14 Gy) to compare the effects of FLASH and CONV irradiation on intestinal epithelial integrity and function. Radiation-induced intestinal injury develops when the intestinal epithelium is damaged resulting in fluid and electrolyte loss as well as bacterial translocation. Similar to human patients, mice exposed to high doses of radiation experience abnormal stool production as a result of damage to the intestinal epithelium. By day 3 post-irradiation, both 14 Gy FLASH and CONV irradiated mice exhibited a significant decrease in the number of formed stool pellets (**Figure 2A-B**). However, the FLASH treated mice had a lesser decrement and faster recovery of stool formation and body weight compared to CONV treated mice **(Figure 2A-B, Supplementary Figure 2A-B)**. By day 6, the CONV treated mice had 22.6% stool production whereas the FLASH treated mice had 47.5% stool production compared to unirradiated controls **(Figure 2B)**. We next compared epithelial integrity following FLASH and CONV irradiation using the FITC-dextran assay where FITC conjugated dextran is fed to the mice and the level of FITC-dextran in the serum reflects the permeability, or loss of barrier function, of the intestinal epithelium ^14^. Mice treated with 14 Gy abdominal CONV irradiation had an increase in FITC-dextran within the serum at 96 hours post-irradiation whereas FLASH treated mice had levels comparable to unirradiated control mice **(Figure 2C)**. Consistent with a functional sparing of the intestine from radiation induced toxicity, we observed a 2.4-fold higher number of regenerating crypts in the jejunum of 14 Gy FLASH treated mice compared to 14 Gy CONV treated mice (**Figure 2D-E**). Notably, an increase in regenerating crypts was also observed in 12 Gy FLASH treated mice compared to 12 Gy CONV treated mice **(Supplementary Figure 2C-D).** These findings demonstrate that FLASH irradiation produces less radiation-induced intestinal injury following abdominal irradiation in healthy mice.

**Figure 2:**
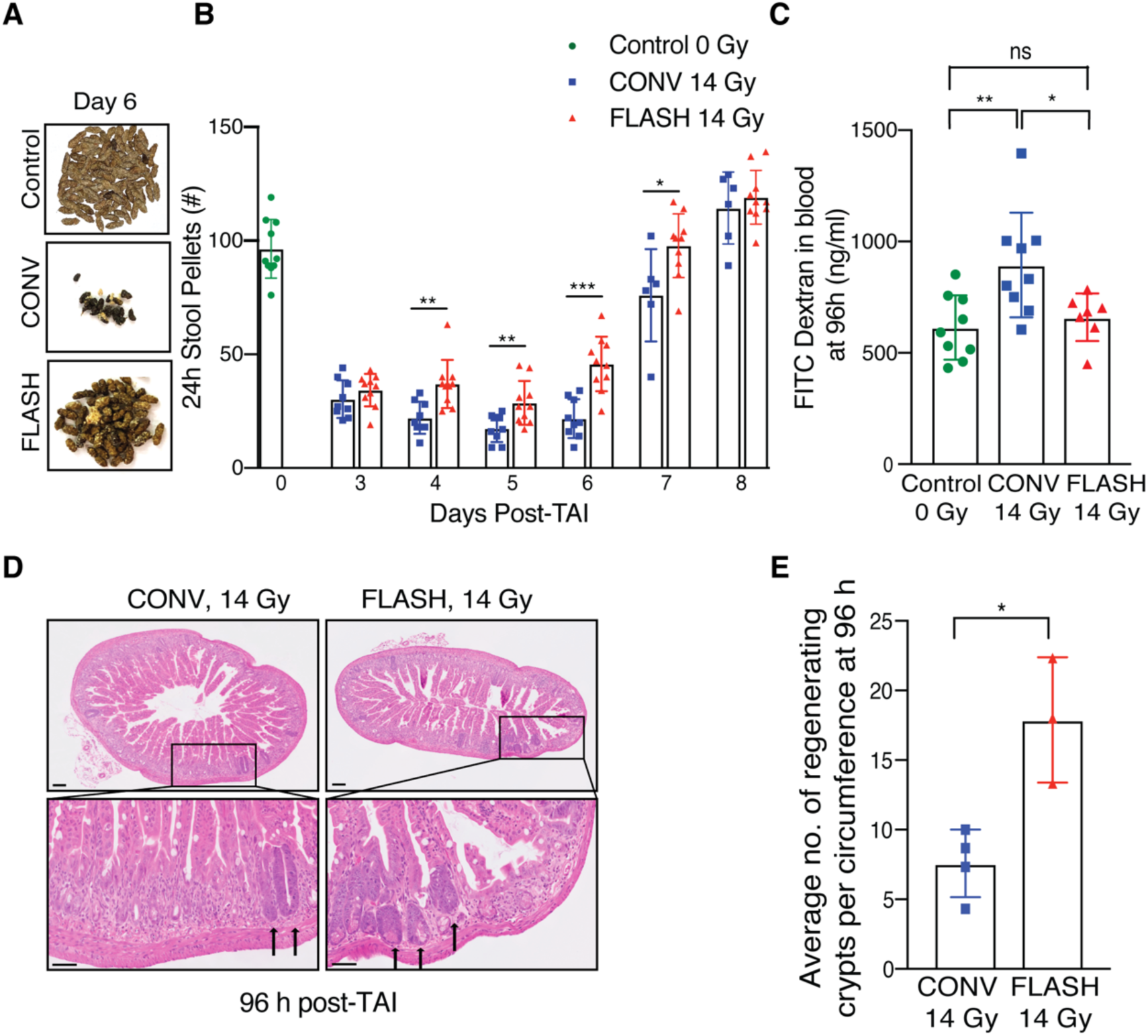
FLASH spares intestinal function and epithelial integrity compared to CONV irradiation at a sub-lethal dose in non-tumor bearing mice. **(A)** Representative images of formed stool pellets from singly housed unirradiated control mice and mice six days after sub-lethal 14 Gy TAI, showing an increase in stools after FLASH vs. CONV. **(B)** Quantification of the number of formed stool pellets excreted over 24 h at the indicated time points from unirradiated control mice and mice after 14 Gy TAI, demonstrating less severe decrement and faster recovery after FLASH vs. CONV irradiation (n=10 mice in unirradiated control & FLASH; n=9 in CONV; n=6 in CONV on days 7 and 8). **(C)** Quantification of FITC-dextran in serum at 96 h after 14 Gy TAI, demonstrating loss of intestinal barrier function after CONV but not FLASH irradiation (n=9 mice in unirradiated control & CONV; n=7 in FLASH). **(D)** Histological images of H&E stained jejunal sections from animals 96 h after 14 Gy TAI. Arrows point to regenerating crypts. **(E)** Quantification of the average number of regenerating crypts per jejunal circumference 96 h after 14 Gy TAI, demonstrating over double the number of regenerating crypts after FLASH vs. CONV irradiation (n=4 mice in CONV; n=3 in FLASH; 3 circumferences per mouse were analyzed). *p<0.05, **p<0.01, ***p<0.001. CONV vs. FLASH compared by unpaired 2-tailed Student’s t-test. Unirradiated control vs. CONV or FLASH compared by one-way ANOVA followed by Tukey’s multiple comparisons post-hoc test. Scale bar: 100 μm. Error bars represent standard deviation of the mean.

### FLASH irradiation alters the proliferation kinetics in crypt cell regeneration compared to CONV

To further investigate the effect of FLASH RT on crypt regeneration and to track the crypt proliferation kinetics, we pulse labelled the irradiated animals with BrdU 2 hours prior to sacrifice. BrdU pulse labelling captures the response of crypt cells entering S-phase and undergoing DNA replication. It is known that radiation-induced DNA damage rapidly induces the G1/S checkpoint through ATM and p53 dependent mechanisms followed by the emergence of regenerating BrdU+ crypts at 96 hours post-irradiation ^15^. As previously reported, we observed a decrease in the number of BrdU+ cells per crypt from 4-72 hours post-irradiation, followed by the appearance of BrdU+ regenerated crypts at 96 hours in the CONV treated group (**Figure 3A-C, Supplementary Figure 3A**, ^16^). While the FLASH treated mice also had a decrease in BrdU+ crypt cells from 4-48 hours, BrdU+ regenerated crypts began to appear as early as 72 hours post-irradiation resulting in a significant increase in BrdU+ cells per crypt at both 72 and 96 hour timepoints post-irradiation (**Figure 3A-C**). These findings further suggest that crypt regeneration is more robust after FLASH compared to CONV.

**Figure 3:**
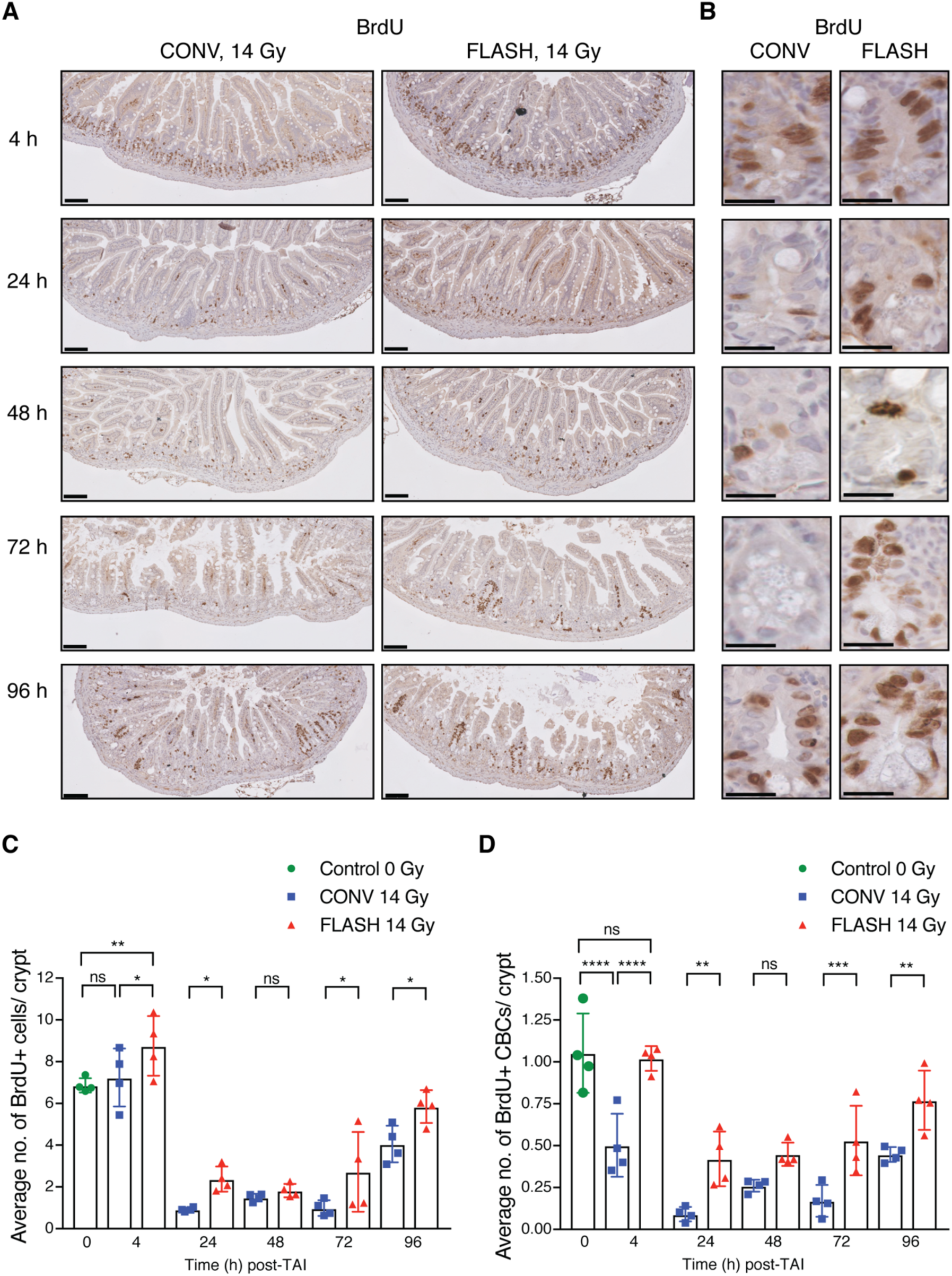
FLASH alters the proliferation kinetics in crypt cell regeneration compared to CONV irradiation in non-tumor bearing mice. Cross sections of small intestines were analyzed for BrdU incorporation and crypt cell proliferation kinetics by BrdU IHC. **(A)** Representative images of BrdU stained jejunum cross sections at 10X magnification at the indicated time points after sublethal 14 Gy TAI. Scale bar: 100 μm. **(B)** 40X magnified crypt images of the jejunum showing BrdU+ crypt cells/CBCs at the indicated time points in CONV and FLASH irradiated animals. Scale bar: 25 μm **(C)** Quantification of the average number of BrdU+ cells per crypt and **(D)** BrdU+ CBCs per crypt at 4 h, 24 h, 48 h, 72 h and 96 h in unirradiated controls and after 14 Gy TAI, demonstrating relative early sparing of proliferating CBCs and more robust regeneration after FLASH *vs*. CONV irradiation. 50 crypts per circumference were quantified for BrdU+ crypt cells and BrdU+ CBCs. 3 circumferences per mouse were analyzed. n=4 mice per group. *p<0.05, **p<0.01, ****p<0.0001; Comparisons by ordinary one-way ANOVA followed by Sidak’s multiple comparisons test. Error bars represent standard deviation of the mean.

Crypt regeneration following radiation injury is mediated by intestinal stem cells located at the base of the crypt. Crypt base columnar (CBC) cells are located at the +1 to +3 position wedged between Paneth cells (**Supplementary Figure 3B**, ^17^). These cells express the leucine-rich repeat-containing G protein-coupled receptor (Lgr5) and are continuously cycling to maintain intestinal homeostasis ^18^. Due to the proliferative nature of Lgr5+ stem cells, they are depleted upon exposure to high dose radiation ^19^. Lineage tracing studies have demonstrated that another Bmi+ quiescent and radioresistant intestinal stem cell population, located in the +4 to +6 region above Paneth cells, compensates for the loss of Lgr5+ cells and gives rise to Lgr5+ cells following injury ^19,20^. We therefore sought to investigate whether FLASH alters the proliferation of the CBC stem cell compartment. Interestingly, we found that in comparison to the overall crypt cell population, the number of BrdU+ intestinal stem cells in the CBC compartment was reduced by 52% in the CONV treated mice at 4 hours, whereas the number of BrdU+ CBCs in the FLASH treated mice were similar to unirradiated control mice (**Figure 3D**). By 24 hours, the percentage of BrdU+ CBCs dropped from 100% to 8% in the CONV treated and to 40% in the FLASH treated mice (**Figure 3D**). By 96 hours post-irradiation, 43% and 73% of CBCs were BrdU+ in the CONV and FLASH treated mice, respectively, correlating with the appearance of regenerating crypts at 96 hours post-irradiation (**Figure 3A-D, Figure 2D**). These data demonstrate that FLASH irradiation preserves the proliferation of CBCs particularly at early timepoints following irradiation in comparison to CONV irradiation.

### FLASH irradiation produces less apoptosis in crypt base columnar cells and modestly less early DNA damage than CONV

The increase in regenerating crypts in FLASH compared to CONV treated mice indicates that intestinal stem cells that repopulate the damaged crypts may be spared from cell death following FLASH. Therefore, we analyzed crypt cell apoptosis by TUNEL and cleaved caspase-3 staining at 4 and 24 h post-irradiation. Within the entire crypt cell population, there were no significant differences in TUNEL positive or cleaved caspase-3 positive cells between FLASH and CONV irradiation at 4 and 24 h **(Supplementary Figure 4A-B)**. In contrast, the number of TUNEL positive and cleaved caspase-3 positive CBCs were decreased in FLASH compared to CONV treated mice **(Figure 4A-D)**. These data indicate that FLASH irradiation produces less apoptosis in intestinal CBCs compared to CONV.

**Figure 4:**
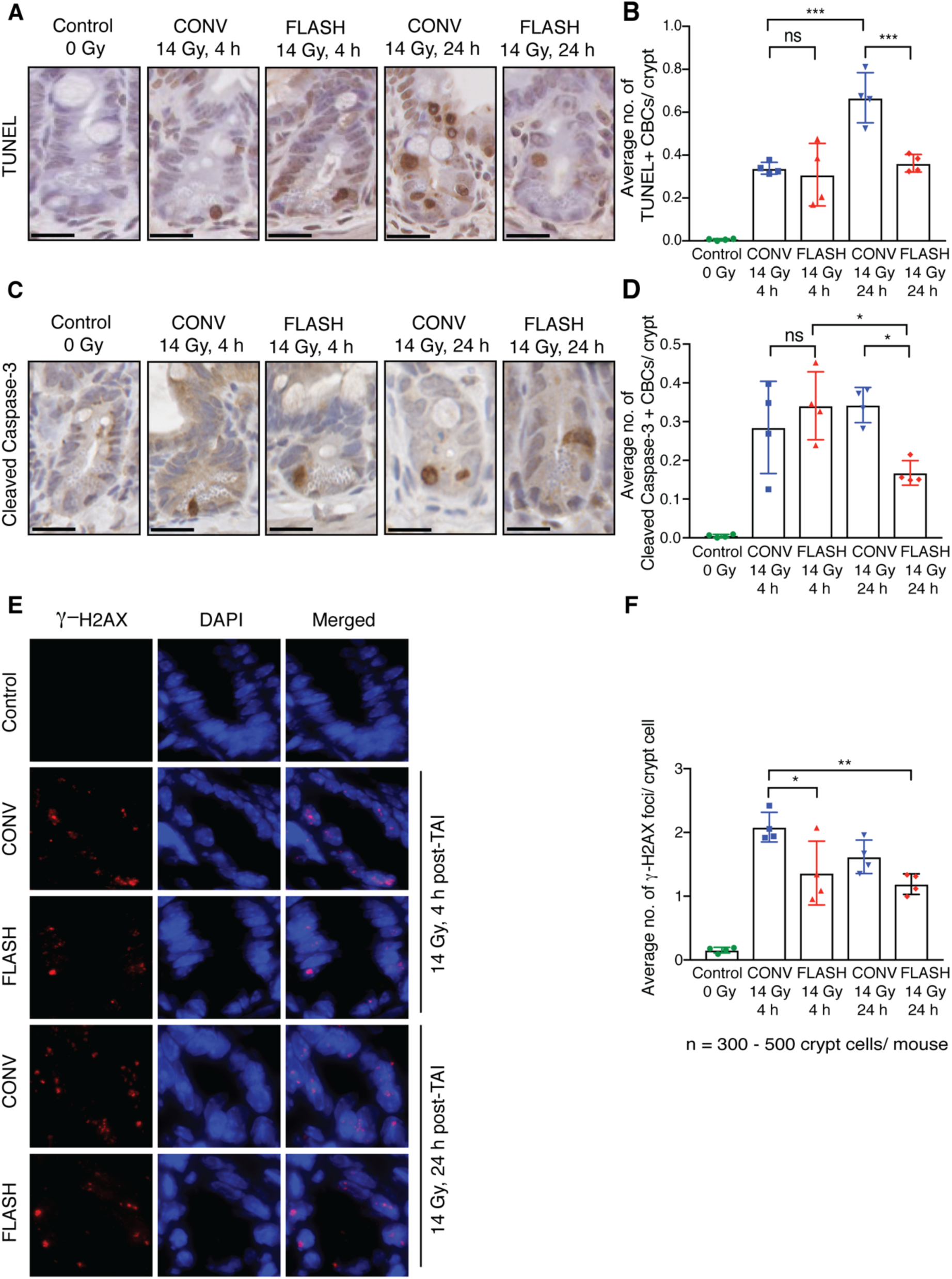
FLASH produces less apoptosis in crypt base columnar cells (CBCs) and less DNA damage than CONV irradiation in non-tumor bearing mice. The jejunum was harvested from unirradiated control mice or at the indicated time points after sublethal 14 Gy TAI and paraffin embedded cross-sections were stained for TUNEL, cleaved caspase-3 and γ-H2AX. **(A)** Representative TUNEL stained images of crypts at 40X magnification and **(B)** quantification of the average number of TUNEL+ CBCs per crypt in the jejunum at 4 h and 24 h after 14 Gy TAI. **(C)** Representative cleaved caspase-3 IHC images of crypts at 40X magnification and **(D)** quantification of the average number of cleaved caspase-3+ CBCs per crypt in the jejunum at 4 h and 24 h after 14 Gy TAI. Scale bar: 25 μm. TUNEL+ CBCs and cleaved caspase-3+ CBCs were quantified per crypt. Crypts in 3 circumferences per mouse were analyzed. n=4 mice per group. Apoptosis in CBCs by TUNEL and cleaved caspase-3 was less at 24 hours after FLASH *vs*. CONV irradiation. **(E)** Representative images of crypts at 100X magnification and **(F)** quantification of γ-H2AX immunofluorescent foci in the crypt cells of jejunum at 4 h and 24 h after 14 Gy TAI, demonstrating fewer foci indicating less early DNA damage and/or increased repair at 4 h after FLASH *vs*. CONV irradiation. n=4 mice per group. An average of 300-500 crypt cells per mouse from 3 circumferences were quantified for γ-H2AX foci. Blue, DAPI; Red, γ-H2AX. ns=no significant difference, *p<0.05, **p<0.01, ***p<0.001; Unirradiated control versus CONV or FLASH compared by one-way ANOVA followed by Tukey’s multiple comparisons test. Error bars represent standard deviation of the mean.

To begin to investigate the molecular mechanisms by which FLASH irradiation spares radiation-induced cell death, we quantified the number of γ-H2AX positive DNA strand breaks in the intestinal crypt cells of mice treated with 14 Gy FLASH or CONV irradiation. The number of γ-H2AX positive DNA breaks (foci) per crypt cell in the jejunum of FLASH treated mice was 1.5 fold less compared to the CONV treated mice at 4 hours post irradiation (**Figure 4E-F)**. By 24 hours, the number of γ-H2AX foci were comparable between FLASH and CONV treated mice **(Figure 4F)**. These findings indicate there is a modest decrease in the initial DNA strand breaks and/or increase in DNA repair in intestinal crypt cells following FLASH irradiation.

### FLASH RT has similar tumor control efficacy as CONV RT and produces less intestinal injury in a preclinical mouse model of ovarian cancer peritoneal metastasis

The findings above demonstrate that FLASH irradiation produces less intestinal injury, raising the intriguing possibility that FLASH irradiation may be an effective strategy to enhance the therapeutic index of radiotherapy for abdominal and pelvic tumor disease. Therefore, we compared the efficacy of FLASH and CONV RT in a preclinical model of ovarian cancer metastasis. We chose to study the ID8 ovarian cancer peritoneal metastasis model in C57BL/6 mice as the disease metastasizes to the peritoneal cavity and significantly affects the bowel, forming tumor nodules along the small and large intestine ^21^. Moreover, radiation-induced bowel toxicity limits the use of radiation therapy in the treatment of ovarian cancer ^22^. In this preclinical study, C57BL/6 mice were injected i.p. with ID8 ovarian cancer cells. At day 10 after injection, the mice were randomized into unirradiated control, 14 Gy CONV, or 14 Gy FLASH irradiation treatment groups (**Figure 5A**). Ninety-six hours post-irradiation, mice were separated and singly housed to quantify stool production. At day 31, when the unirradiated control mice display signs of morbidity, all mice were euthanized, and tumor burden was quantified (**Figure 5A**). FLASH treated mice lost only 10% of their body weight while the CONV mice lost 25% of their body weight 6 days post-TAI, all of which restored their pre-irradiation bodyweights within 2 weeks after irradiation **(Supplemental Figure S5A)**. Moreover, the number of stool pellets excreted over 24 hours at day 5 post-irradiation and their corresponding stool weights were higher in FLASH compared to CONV treated mice **(Figure 5B, Supplemental Figure S5B)**. These data indicate that FLASH RT produces less intestinal injury in tumor-bearing mice compared to CONV RT. Analysis of total tumor burden revealed a robust decrease in both the number of tumor nodules and total tumor weights in mice treated with both CONV and FLASH irradiation when compared to the unirradiated control mice **(Figure 5C-E)**. No significant differences in tumor number or weights were observed when comparing FLASH to CONV treated mice, indicating that FLASH and CONV irradiation have similar efficacy in the treatment of ovarian cancer peritoneal tumors (**Figure 5D-E**). Our results demonstrate that while both FLASH and CONV RT reduced the ovarian cancer tumor burden in the peritoneal cavity, FLASH RT also produced less radiation-induced intestinal injury in this tumor model.

**Figure 5:**
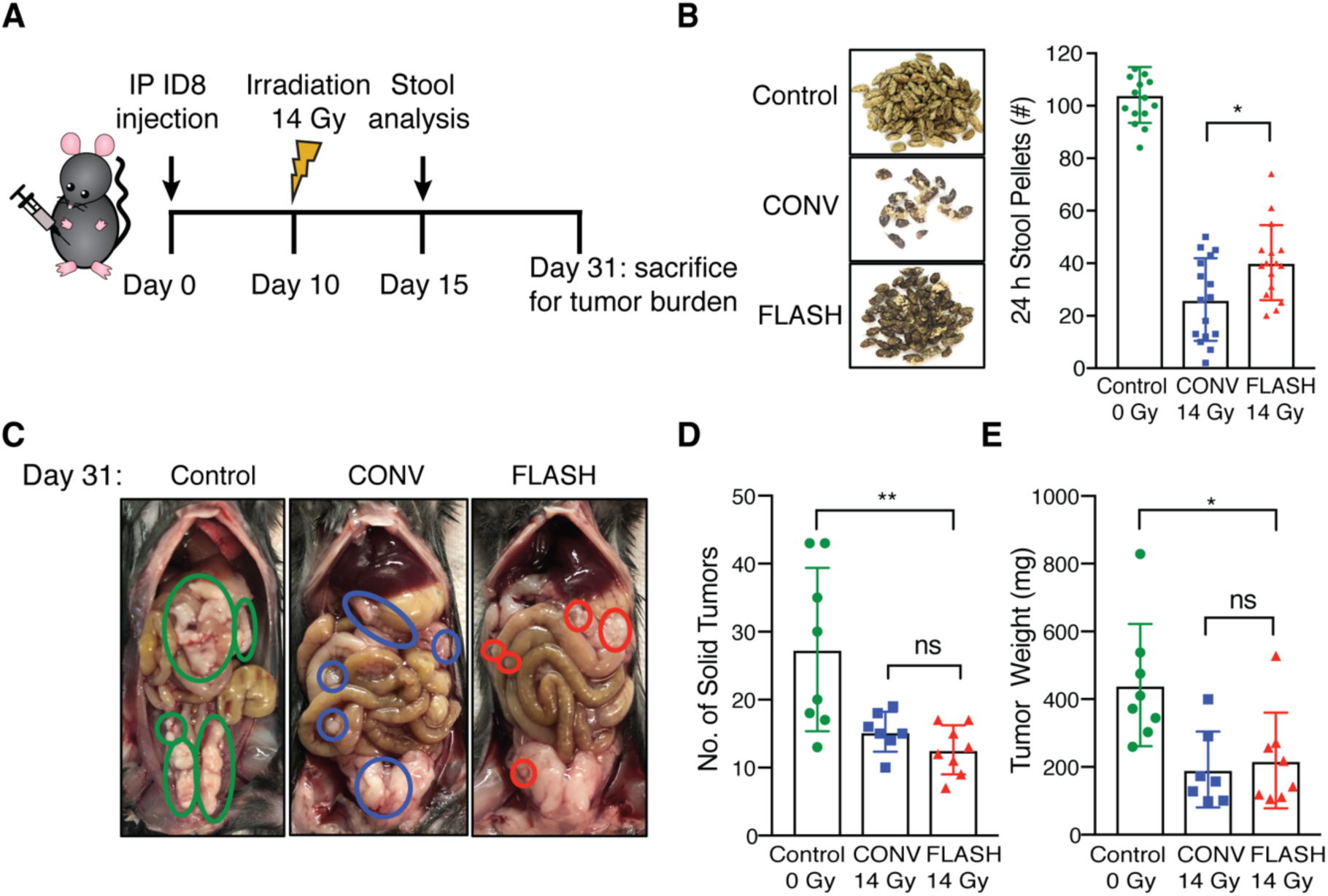
FLASH has similar tumor control efficacy as CONV irradiation and produces less intestinal injury in a preclinical syngeneic ovarian cancer mouse model. **(A)** Schematic of the ID8 tumor cell peritoneal injection, total abdominal irradiation, assessment of intestinal function by stool counts and weights, euthanasia and tumor burden analysis at the indicated time points in 6-8 week old female C57BL/6 mice. **(B)** Representative images of stool pellets and quantification of the number of stool pellets excreted over 24 h at 15 days after tumor cell injection, or 5 days after sublethal 14 Gy TAI (n=16 mice per group), showing increased stools after FLASH *vs*. CONV, as also seen in non-tumor bearing mice. **(C)** Representative images displaying metastatic tumor burden macroscopically in mice that were intraperitoneally injected with ID8 ovarian tumor cells and **(D and E)** quantification of the number of solid tumor nodules and tumor weights in unirradiated control mice and mice treated with 14 Gy CONV or FLASH irradiation, demonstrating similar tumor control efficacy with both FLASH and CONV irradiation. n=8 mice per group; ns=no significant difference, *p<0.05, **p<0.01; Comparisons by ordinary one-way ANOVA followed by Tukey’s multiple comparisons test. Error bars represent standard deviation of the mean.

## Discussion

Epithelial ovarian cancer (EOC) is the 5^th^ leading cause of cancer related deaths among women in the United States. The majority of women diagnosed with epithelial ovarian cancer present with advanced stage disease and widespread metastasis throughout the abdomen and pelvis. First-line therapies for patients with EOC include optimal surgical debulking and platinum-based chemotherapy. While the majority of patients initially respond to platinum-based chemotherapy, most develop recurrent disease and succumb to their disease. With the development of chemoresistance in the majority of patients, the identification of effective therapeutic strategies for the treatment of ovarian cancer is needed.

Our findings have important clinical implications for the treatment of advanced EOC. We demonstrate that both FLASH and CONV irradiation have similar tumor control efficacy in a preclinical model of ovarian cancer metastasis. Importantly, we demonstrate that abdominal FLASH RT spares mice from lethal intestinal injury relative to CONV RT. Moreover, at sublethal doses, FLASH RT preserves intestinal function and recovery compared to CONV RT in both healthy and tumor-bearing mice. These observations of marked radiobiological advantages of FLASH over conventional dose rate radiotherapy in the highly radiosensitive intestine suggest the translational potential of FLASH RT for increased therapeutic index in the treatment of metastatic ovarian cancer. While the preclinical technologies that have been used for FLASH irradiation of mice do not scale to larger, deep-seated tumors in human patients, new radiation therapy technologies to translate FLASH RT to humans are actively under development ^23^. Additionally, studies to determine whether FLASH mediated protection of the intestine from radiation-induced toxicity is maintained in fractionated protocols will also be needed prior to clinical trials.

The mechanisms of FLASH sparing of the intestine likely stem from the enhanced regenerative potential of intestinal stem cells. Radiation causes damage to the intestinal epithelium by inducing cell death in Lgr5+ intestinal crypt stem cells that regenerate the intestinal epithelial cells ^17^. We observed that at large single doses of 12-16 Gy, with responses spanning non-lethal to completely lethal intestinal injury with conventional dose rate irradiation, FLASH RT produced modestly fewer γ-H2AX foci within intestinal crypt cells and less apoptosis in CBC stem cells, a greater number of regenerating crypts, and preservation of intestinal function and survival. These findings suggest that FLASH RT produces less DNA damage and/or alters the DNA damage response in intestinal stem cells to enhance crypt regeneration. Our findings set the stage for future studies investigating the cellular and molecular mechanisms driving the FLASH response in the intestinal stem cell compartment.

Previous studies have suggested that altered oxygen radiochemistry at extremely high dose rates may underlie the FLASH effect *in vitro* and *in vivo* ^24,25^. Based on radiochemical modeling studies, a 10 Gy or greater dose of FLASH RT has the potential to deplete molecular oxygen in tissues at physiologic oxygen tensions, as oxygen rapidly reacts with radicals formed by radiolysis of water and other biomolecules (ROS) ^26^. Perhaps especially in stem cell niches that may have baseline hypoxia, transient anoxia produced by FLASH but not conventional dose rate irradiation would result in reduced DNA double strand breaks caused by reduced oxygen-mediated fixation of DNA damage ^27,28^. In support of this hypothesis, Montay-Gruel *et al.* found that carbogen breathing to increase tissue oxygenation partially abrogated the cognitive sparing observed after FLASH brain irradiation ^29^. We demonstrate that FLASH RT produces less early DNA damage within intestinal crypt cells *in vivo* compared to CONV RT, a finding consistent with the oxygen depletion hypothesis. Future studies to measure tissue oxygen and ROS levels in real time during FLASH and CONV irradiation, which may be enabled by oxygen and ROS sensing nanoparticle probes, would provide the most direct evidence of this mechanism ^30,31^.

In summary, our findings suggest that with continued development, FLASH irradiation may offer an opportunity to reintroduce radiotherapy into the armamentarium of ovarian cancer therapeutics. Historical data demonstrates that the majority of ovarian tumors respond to radiotherapy. Therefore, FLASH irradiation has the potential to extend overall survival and improve quality of life for women diagnosed with advanced epithelial ovarian cancer.

## Materials and Methods

### Study design

The goal of this study was to investigate the safety and efficacy of total abdominal FLASH RT (TAI-FLASH) compared to conventional dose rate RT (TAI-CONV) in reducing ovarian cancer peritoneal metastases. All procedures for use of animals and their care were approved by the Institutional Animal Care and Use Committee of Stanford University in accordance with institutional and NIH guidelines. Six to eight-week-old female C57BL/6 mice (Jackson Labs) were irradiated by both CONV and FLASH methods. Mice were anaesthetized with a mixture of ketamine (100 mg/kg) and xylazine (10 mg/kg) injected into the peritoneum. Control mice were treated with the same dose of ketamine/xylazine but were not exposed to radiation. Safety of TAI-FLASH relative to TAI-CONV was first assessed in non-tumor bearing mice. At a lethal dose of 16Gy (when delivered by CONV), mice were irradiated by FLASH and CONV methods and the overall survival and the number of regenerating crypts within the jejunum were determined. The effects of FLASH and CONV irradiation on gastrointestinal function and epithelial integrity were evaluated at a sublethal dose of 14Gy. GI function and integrity were measured by the number of stool pellets excreted over 24 h, regenerating crypts and FITC dextran assay 96 h post-irradiation. Mechanisms behind the normal tissue-sparing effect of FLASH irradiation was investigated by studying crypt regeneration, crypt cell proliferation kinetics at 4 h, 24 h, 48 h, 72 h and 96 h after irradiation, crypt cell damage and apoptotic crypt cell death kinetics at 4 h and 24 h after irradiation. The efficacy of abdominal FLASH RT and CONV RT was then compared in a preclinical model of ovarian cancer metastasis. ID8 ovarian cancer peritoneal metastasis was established in C57BL/6 mice. Analysis of macroscopic tumor burden and tumor weight as well as GI function as described above was performed to demonstrate tumor control efficacy while sparing normal GI function.

### Mouse irradiation

We developed a custom mouse stereotactic positioning frame made of PLA plastic using 3-D printing. Reproducible positioning within the frame was achieved by registering the front teeth on a nylon filament at a fixed location at the cranial end of the frame with extension of the hindlimbs and tail through designated slots at the caudal end of the frame (Supplementary Figure 1A). Positioning reproducibility to within 1 mm was confirmed by microCT imaging in a representative cohort of mice (Figure 1A). An abdominal irradiation shield was made by 3-D printing a PLA plastic shell with a central opening of 4 cm (lateral) by 3 cm (craniocaudal). Internally, a 3 cm thick layer of aluminum oxide powder in tandem with a 1 cm thick layer of tungsten spheres (2 mm diameter) was placed, a combination designed to minimize leakage dose from bremsstrahlung radiation produced in the shield materials (Supplementary Figure 1A). The stereotactic frame was registered to the shield such that the opening extended from the tenth rib at the cranial border to 3 cm caudally, consistently encompassing all of the small intestine. EBT3 Gafchromic film (Ashland Advanced Materials, Bridgewater NJ) dosimetry confirmed that leakage dose to the shielded portions of the body when irradiating with 16 MeV electrons was <3.5% of the central dose at 10 mm from the field edge and further.

The shield and positioning frame were loaded into a polystyrene cradle registered to specified locations relative to the LINAC treatment head for FLASH and CONV irradiation. For both FLASH and CONV setups, we placed EBT3 Gafchromic films between layers of polystyrene to measure transverse and depth dose profiles to confirm dose homogeneity throughout the treatment volume (Supplementary Figure 1D-F). In addition, entrance dose for every individual mouse irradiation was recorded by EBT3 Gafchromic films (1×2 inch) placed inside the positioning frame.

### CONV irradiation setup

We used a Varian Trilogy radiotherapy system (Varian Medical Systems, Palo Alto, CA) to perform both CONV and FLASH irradiation. For CONV irradiation, the gantry was rotated to 180 degree (beam direction from floor to ceiling) and the collimator was rotated to 0 degree. The cradle with the shield and mouse jig was placed on top of a 15 cm electron applicator, supported by a 1 cm thick lead sheet with a 3.5 × 5 cm opening that also served as a primary shield, such that the distance from the electron scattering foil to the shield was 77.6 cm. Under service mode, irradiation was delivered using a clinical 16 MeV electron beam in the 400 MU/minute dose rate mode (pulse repetition rate 72 Hz). Calibration by film dosimetry determined that the entrance dose after the shield was 1.18 cGy/MU, with a resulting average dose rate of 0.079 Gy/s (dose per pulse of 0.00109 Gy).

### FLASH irradiation setup

We configured the Varian Trilogy radiotherapy system to perform FLASH irradiation as previously described ^32^. The gantry was rotated to 180 degree, the treatment head cover was removed, and the jaws were fully opened (40 × 40 cm). The cradle with radiation shield and mouse stereotactic frame was loaded and registered to fixed points on the face of the gantry, such that the distance from the electron scattering foil to the shield was 14.6 cm. Beam parameters were configured on a dedicated electron beam control board. We used an electron beam energy of approximately 16 MeV with the 16 MeV scattering foil (confirmed by depth dose measurements) and adjusted the radiofrequency power and gun current settings to produce a dose per pulse of 2.0 Gy. We controlled pulse delivery using a programmable controller board (STEMlab 125-14, Red Pitaya, Solkan, Slovenia) and relay circuit to count the number of delivered pulses detected by the internal monitor chamber and impose beam hold and release through the respiratory gating system of the LINAC. We used an external ion chamber positioned after the mouse and 10 cm of solid water (where the dose rate did not saturate the chamber), calibrated to film measurements of entrance dose, to provide immediate dose readout per mouse. The pulse repetition rate was set to 108 Hz for an average dose rate of 216 Gy/sec at 2 Gy/pulse.

### Ovarian cancer tumor model

The mouse ID8 ovarian cancer cell line was obtained from Dr. Katherine F. Roby, University of Kansas Medical Center, Kansas City, KS ^21^. The cell line was authenticated from the original source and was used within 6 months of receipt. Additionally, cells were tested upon receipt for viability, cell morphology and the presence of *Mycoplasma* and viruses (Charles River Laboratories). ID8 cells were passaged in DMEM supplemented with 10% FCS and Pen/Strep. Cells (5×10^6^ cells in 200 μl PBS) were administered via intraperitoneal injection using a 27-gauge needle (BD). Injected animals were treated with FLASH or CONV RT on day 10 post-inoculation. Animals were euthanized on day 31 and tumor burden was assessed.

### Survival and intestinal function analysis

Survival times of mice were measured from irradiation until death or morbidity requiring euthanasia. Global gastrointestinal function was assessed by measuring daily weights.

Intestinal function was also assessed by counting the number of formed stool pellets at specified time points, as diarrhea and inability to make stools is an indicator of radiation-induced gastrointestinal syndrome. Animals were singly housed for 24 h prior to stool collection, after which their droppings were separated from the cage bedding and counted. Animals were then returned to their original cage groupings.

Intestinal barrier function was assessed in a cohort of mice by the FITC-dextran assay. FITC-dextran (Sigma-Aldrich) was prepared as a stock solution at a concentration of 100 mg/mL in PBS. FITC-dextran was administered via oral gavage at a dose of 0.6 mg FITC-dextran/g of body weight at 96 hours post-irradiation. Mice were sacrificed after 4 h and blood was collected by cardiac puncture. After incubating 2 h at room temperature, serum was isolated by centrifugation at 2,000 X g for 20 minutes at 4°C. The amount of FITC-dextran in the serum was determined by measuring the fluorescence at an excitation wavelength of 485 nm and an emission wavelength of 525 nm using a plate reader (Synergy microplate reader, H1 Biotek).

### Tissue processing and histological analysis

Animals were euthanized via CO2 asphyxiation and secondary cardiac exsanguination. Soft tissues were harvested and immersion-fixed in 10% neutral buffered formalin for 24 h, followed by PBS for 24 h and then stored in 70% ethanol. Three transverse sections of the small intestine from the jejunum (mid-segment) were collected. Formalin-fixed tissues were processed routinely, embedded in paraffin, sectioned at 5 μm and stained with hematoxylin and eosin.

### Regenerating crypt counts

A total of three transverse sections of jejunum were analyzed per mouse for the number of regenerating crypts by the crypt microcolony assay ^33^. Transverse sections were analyzed if they met the following criteria: 1) a complete jejunal circumference was present; and 2) the mucosa was oriented perpendicular to the long axis of the intestine. Crypts were considered regenerating if they comprised >10 basophilic crypt epithelial cells (n=4 mice/ group).

### γ-H2AX immunofluorescence

Slides were baked and deparaffinized in xylene and alcohol series followed by heat-mediated antigen retrieval using sodium citrate buffer. The tissue sections were then blocked with serum-free protein blocking agent (Dako) and incubated with rabbit mAb for Phospho-Histone H2A.X (Ser139) (20E3) (Cell Signaling Technology) at 4°C overnight. Slides were washed in 1X PBS and incubated with Alexafluor 594 conjugated anti-rabbit secondary antibody for 30 min at 37°C. 1X PBS containing 0.1% BSA and 0.2% Triton X-100 were used to dilute the antibodies. The slides were then washed and quenched for autofluorescence using Sudan Black B for 10 min at room temperature followed by DAPI and mounting. Images were captured as Z-stacks using a fluorescence microscope (Leica DMi8). Maximum intensity projections of the captured Z-stacks were analyzed for γ-H2AX foci. An average of 300-500 crypt cells per mouse were analyzed for the number of punctate immunoreactive γ-H2AX foci in 3 jejunal circumferences per mouse (n=4 mice/ group).

### TUNEL assay

TUNEL assay was performed using ApopTag® Peroxidase In Situ Apoptosis Detection Kit (Millipore) according to manufacturer’s instructions. Briefly, deparaffinized sections were enzymatically digested using 20 μg/ml Proteinase K and quenched in hydrogen peroxide before treating with TdT enzyme and anti-digoxigenin peroxidase conjugate. Signals were developed using DAB peroxidase substrate and signal development was monitored under a light microscope. Slides were scanned using Nanozoomer (Hamamatsu). Number of TUNEL+ crypt cells and TUNEL+ crypt base columnar cells (CBCs) were quantified per crypt per circumference. Cells were considered positive if they exhibited dark brown nuclear staining. Three jejunal circumferences per mouse were analyzed (n=4 mice/ group).

### Cleaved caspase-3 immunohistochemistry

Antigen retrieval was performed in 10 mM sodium citrate buffer. Following avidin/biotin block and serum-free protein block, sections were incubated in cleaved caspase-3 (Asp175) antibody (1:300; Cell Signaling Technology) at 4°C overnight in a humidified chamber. Biotinylated secondary antibody, ABC kit and DAB substrate were employed to develop the signal. Slides were scanned using Nanozoomer. Number of cleaved caspase-3+ crypt cells and cleaved caspase-3+ CBCs were quantified per crypt per circumference. Three jejunal circumferences per mouse were analyzed (n=4 mice/group).

### BrdU immunohistochemistry

BrdU (Sigma) was intraperitoneally injected two hours prior to sacrificing the animals at a dose of 100 mg/kg in sterile PBS. Tissues were serially harvested at their respective timepoints following irradiation, fixed, paraffin-embedded and sectioned. Paraffin sections (5 μm) were baked, deparaffinized and subjected to heat-mediated antigen retrieval in Tris-EDTA pH 9.0. Antigen retrieval was performed at high pressure for 5 min in a pressure cooker. Slides were cooled on ice, rinsed in water and quenched for endogenous peroxidase in 0.3% H_2_O_2_ at room temperature. Sections were permeabilized in 1X TBS containing 0.025% Triton X-100 (TBS-TX) for 10 min, blocked with TBS-TX containing 10% goat serum and 1% BSA for 30 min at room temperature. Tissue sections were incubated with BrdU mouse monoclonal biotinylated primary antibody (1:100; Invitrogen) at 4°C overnight followed by biotinylated anti-mouse secondary antibody. Signals were amplified using Vecta Elite ABC kit and monitored under a light microscope using DAB peroxidase substrate kit. Sections were counter-stained with hematoxylin, dehydrated, and mounted. Slides were scanned using NanoZoomer. Fifty crypts per circumference were quantified for BrdU+ crypt cells and BrdU+ CBCs. Three jejunal circumferences per mouse were analyzed (n=4 mice/group).

### Complete blood count analysis

Blood was collected by cardiac puncture at the time of sacrifice. Complete blood counts data were collected by analyzing the blood using a Hemavet 950FS (Drew Scientific).

### Statistical Analysis

Analysis of two groups with a continuous outcome were performed using a Student’s t-test. Analyses of three groups or more with a continuous outcome were performed using an ANOVA and pair-wise comparisons were performed in a post hoc analysis with a Tukey or Sidak adjustment for multiple comparisons. Error bars represent standard deviations. Time to event outcomes were summarized with Kaplan-Meier curves and groups were compared in a log-rank test. Analyses of continuous outcomes measured at multiple time points were performed using a mixed effects model with pair-wise comparisons done in a post hoc test with a Tukey adjustment for multiple comparisons. All statistical tests were two-sided with an alpha level of 0.05. All analyses were performed in SAS v 9.4 (SAS Institute Inc., Cary, NC) or Prism v 8.3.0 (GraphPad Software, San Diego, CA).

## Funding

This work was supported by the Office of the Assistant Secretary of Defense for Health Affairs through the Department of Defense Ovarian Cancer Research Program under Award No. W81XWH-17-1-0042; the My Blue Dots fund; the Stanford University Department of Radiation Oncology; the Weston Havens Foundation; the Stanford University School of Medicine; the Stanford University Office of the Provost; the Wallace H. Coulter Foundation; the Cancer League; the Swedish Childhood Cancer Foundation; the Foundation BLANCEFLOR Boncompagni Ludovisi n’ee Bildt; the American Association for Cancer Research.

The authors would like to thank Miguel Jimenez, Daniel Pawlak, and James Clayton from Varian Medical Systems for their technical assistance on the FLASH irradiation system.

## Author Contributions

EBR, BWL, PGM, EEG, ACK designed experiments. KL, SN, JW, SC, JTE, PL, RM, FML, ES, MR, RK, AK, DR performed experiments. KL, SN, JW, PM, BWL and EBR analyzed data. LS designed the 3D printed shield for the studies. KB and ASY provided access and support to the clinical linear accelerator. KMC analyzed histological data. RVE provided statistical analysis of the data. EBR, BWL, SN, JW wrote the manuscript. KL, SN, JW, SC, MR, RVE, OD, EEG, KMC edited the manuscript. All authors proofread and approved the manuscript.

## Competing Interests

BWL: Research support from Varian Medical Systems; Board member of TibaRay.

All other authors do not have any competing interest to declare.

