## Supplemental figures 1-5 for "FLASH irradiation enhances the therapeutic index of abdominal radiotherapy for the treatment of ovarian cancer"

### SUPPLEMENTARY FIGURE 1

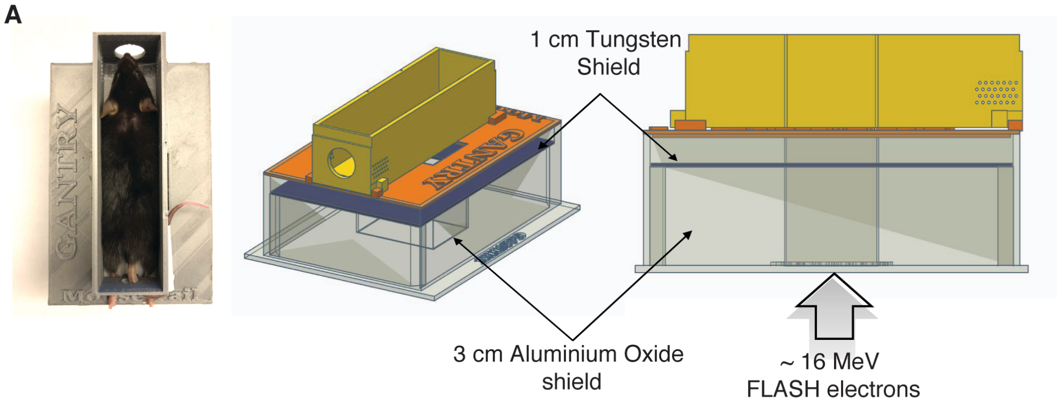


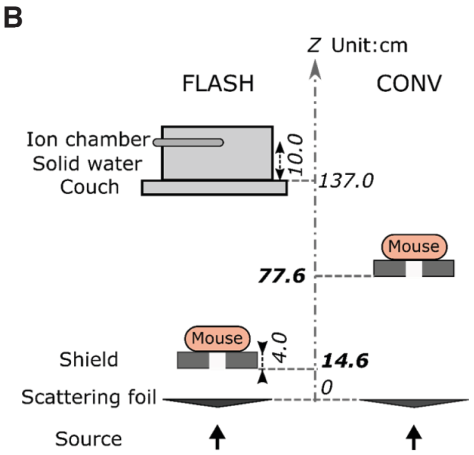


C

| Parameters | FLASH | CONV |
|---|---|---|
| e-beam energy | ~16 MeV | ~16 MeV |
| Repetition rate | 108 Hz | 72 Hz |
| Dose per pulse | 2.0 Gy | 0.00109 Gy |
| Average dose rate | 216 Gy/s | 0.07863 Gy/s |
| Instantaneous dose rate (pulse length 5 $\mu$s) | 4.0E5 Gy/s | 218.5 Gy/s |

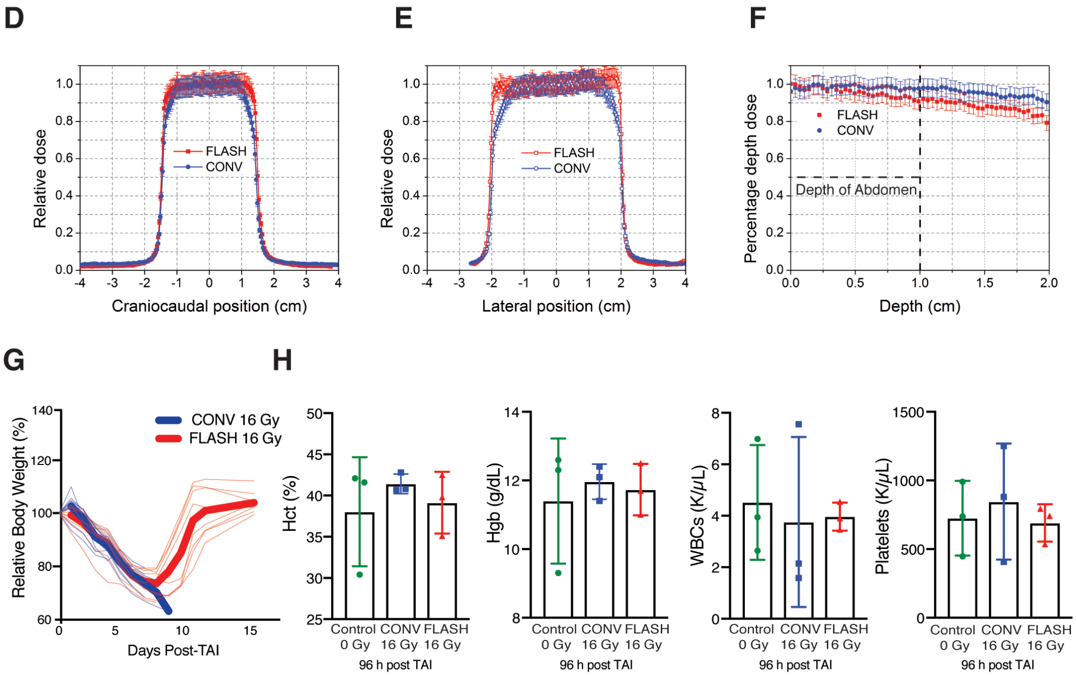

**Supplementary Figure 1: CONV dose rate and FLASH abdominal irradiation set up; and lack of hematologic toxicity after abdominal irradiation in non-tumor bearing mice**

**(A)** Design of 3-D printed PLA plastic mouse stereotactic positioning frame and abdominal irradiation shield comprising a 3-D printed PLA plastic shell containing layers of 3 cm thick aluminum oxide ($Al_2O_3$) powder in tandem with a mixture of 1 cm thick tungsten beads (2 mm diameter) and powder. The $Al_2O_3$ layer slows down the ~16 MeV primary electrons and the tungsten layer stops the transmitted electrons and attenuates the bremsstrahlung x-rays produced by the electron beam. **(B and C)** Schematic of the geometric and beam parameters for FLASH and CONV setups. The shorter distance from the electron scattering foil and higher charge per pulse in FLASH produces an average dose rate 2750 times the CONV dose rate. **(D)** Craniocaudal, **(E)** lateral, and **(F)** depth dose profiles for FLASH and CONV setups. The doses are uniformly distributed within the abdominal region. **(G)** Relative body weight (%) of each mouse over time after 16 Gy CONV or FLASH TAI; bold lines: averages for the cohorts. **(H)** Circulating blood cell counts (hematocrit, hemoglobin, WBCs, and platelets) in unirradiated control mice and irradiated mice 96 h after 16 Gy TAI show no significant hematologic toxicity from either FLASH or CONV abdominal irradiation. Blood was drawn by cardiac puncture at the time of euthanasia. ns=no significant difference by ordinary one-way ANOVA and Tukey's multiple comparisons test. Error bars represent standard deviation of the mean.

#### SUPPLEMENTARY FIGURE 2

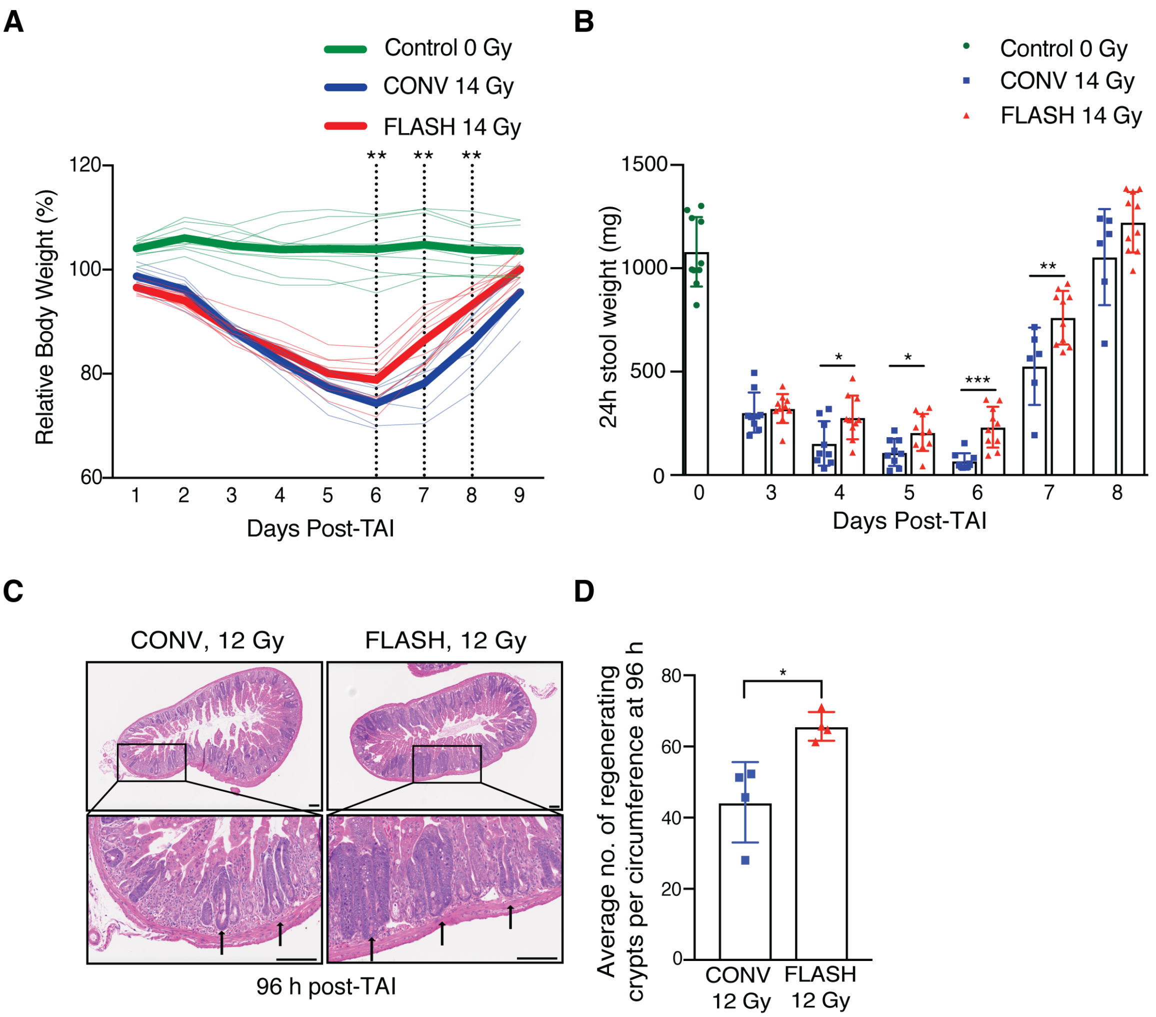


**Supplementary Figure 2: FLASH spares intestinal function and epithelial integrity compared to CONV irradiation at a sub-lethal dose in non-tumor bearing mice**

**(A)** Relative body weight (%) of each mouse over time after 14 Gy CONV or FLASH TAI; bold lines: averages for the cohorts. The stippled lines indicate days at which the difference between CONV and FLASH is significant. (n=10 mice in unirradiated control & FLASH; n=9 in CONV; n=6 in CONV on days 7, 8 and 9). **(B)** Quantification of the weight of formed stool pellets excreted over 24 h at the indicated time points from unirradiated control mice and mice after 14 Gy TAI, demonstrating less severe decrement and faster recovery after FLASH *vs*. CONV irradiation. (n=10 mice in unirradiated control & FLASH; n=9 in CONV; n=6 in CONV on days 7 and 8). **(C)** Histological images of hematoxylin and eosin (H&E) stained jejunal sections from animals 96 h after a lower dose of 12 Gy TAI. Scale bar: 100 μm. Arrows point to regenerating crypts and **(D)** quantification of the average number of regenerating crypts per circumference 96 h after 12 Gy TAI, demonstrating a higher number of regenerating crypts after FLASH *vs*. CONV irradiation. Regenerating crypts were counted in 3 circumferences per mouse. n=4 mice per group. *p<0.05, **p<0.01, ***p<0.001. CONV *vs*. FLASH compared by unpaired 2-tailed Student's t-test. Error bars represent standard deviation of the mean.

**SUPPLEMENTARY FIGURE 3**

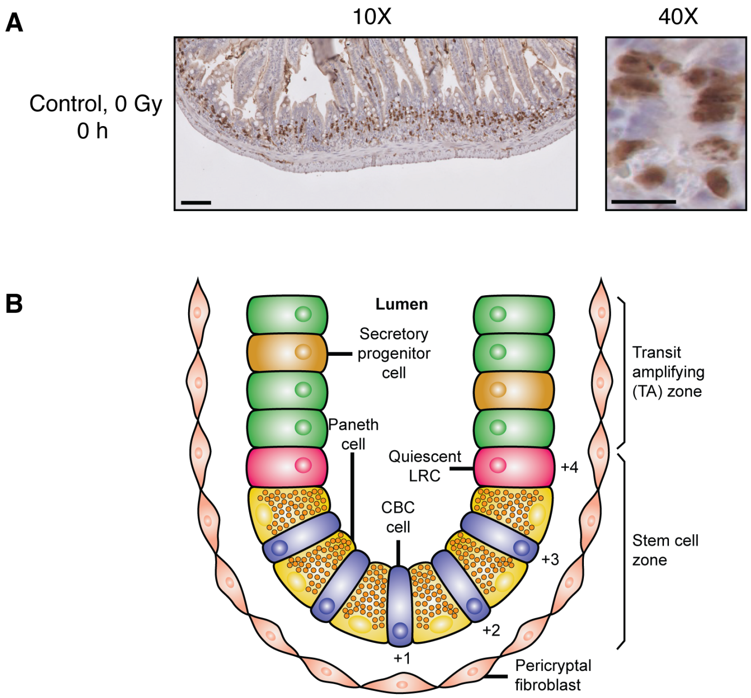


**Supplementary Figure 3: BrdU staining of unirradiated intestinal crypt cells**

**(A)** Representative image of BrdU stained jejunum cross section at 10X magnification and the corresponding 40X magnified image of a crypt showing BrdU+ crypt cells/CBCs from unirradiated control animals. Scale bar: 100 μm at 10X magnification and 25 μm at 40X magnification. **(B)** Schematic representation of the structure of the jejunal crypt with CBCs located at +1, +2 and +3 positions at the base of the crypt and sandwiched between Paneth cells. CBC=Crypt Base Columnar cell; LRC=Label Retaining Cell. The base of the crypt represents the stem cell zone and above this compartment is the transit amplifying (TA) zone. The TA compartment comprises TA cells with secretory progenitor cells interspersed between the TA cells, which matures into the Paneth cell population. Quantification of BrdU+ crypt cells, TUNEL+ crypt cells, and cleaved caspase-3+ crypt cells included cells from both the TA zone and the stem cell zone.

**SUPPLEMENTARY FIGURE 4**

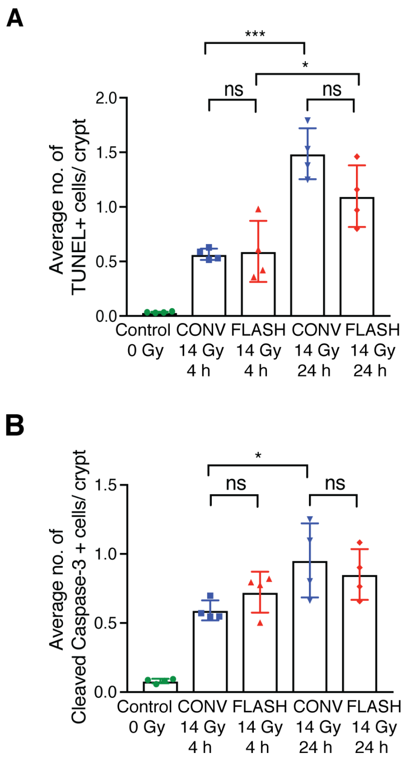


**Supplementary Figure 4: FLASH and CONV irradiation produce similar apoptosis of crypt cells in non-tumor bearing mice**

**(A)** Quantification of the average number of TUNEL+ cells per crypt and **(B)** average number of cleaved caspase-3+ cells per crypt in the jejunum analyzed at 4 h and 24 h after 14 Gy TAI, demonstrating no significant differences in apoptosis between FLASH and CONV irradiation when considering the entire crypt, in contrast to the difference seen when restricting the analysis to only the crypt base columnar cells (CBCs) (Figure 4). TUNEL+ cells and cleaved caspase-3+ cells were quantified per crypt for crypts from 3 circumferences per mouse. n=4 mice per group. ns=no significant difference, *$p<0.05$, ***$p<0.001$. Comparisons by ordinary one-way ANOVA followed by Tukey's multiple comparisons test. Error bars represent standard deviation of the mean.

**SUPPLEMENTARY FIGURE 5**

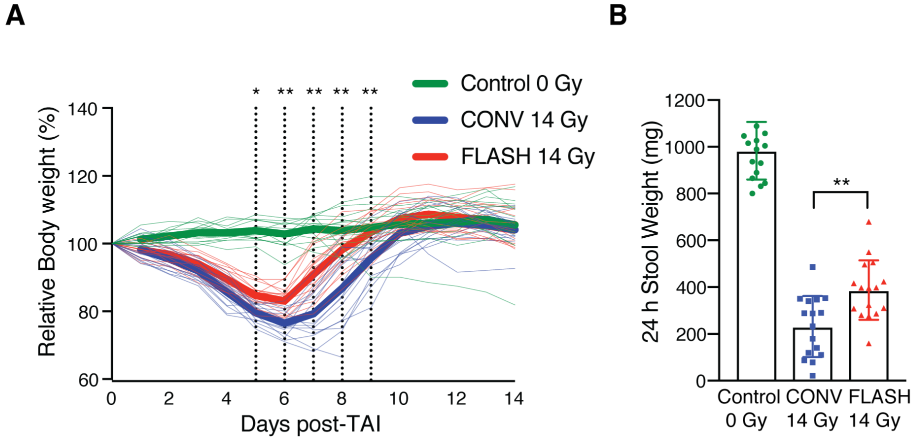


**Supplementary Figure 5: FLASH spares intestinal function compared to CONV dose rate irradiation in a preclinical syngeneic ovarian cancer mouse model**

**(A)** Relative body weight (%) of each ID8 tumor-bearing mouse over time after 14 Gy CONV or FLASH TAI; bold lines: averages for the cohorts. The stippled lines indicate days at which the difference between CONV and FLASH is significant. n=16 per group. **(B)** Quantification of the weights of stool pellets excreted over 24 h at day 5 after 14 Gy TAI by ID8 tumor-bearing mice, showing increased stools after FLASH *vs*. CONV, as also seen in non-tumor bearing mice. n=16 per group. (*$p<0.05$; **$p<0.01$). Comparisons by ordinary one-way ANOVA followed by Tukey's multiple comparisons test. Error bars represent standard deviation of the mean.
